# AXDND1 is required to balance spermatogonial commitment and for sperm tail formation in mice and humans

**DOI:** 10.1101/2023.11.02.565050

**Authors:** Brendan J. Houston, Joseph Nguyen, D. Jo Merriner, Anne E. O’Connor, Alexandra M. Lopes, Liina Nagirnaja, Corinna Friedrich, Sabine Kliesch, Frank Tüttelmann, Kenneth I. Aston, Donald F. Conrad, Robin M. Hobbs, Jessica EM Dunleavy, Moira K. O’Bryan

**Affiliations:** School of BioSciences, Bio21 Institute, The University of Melbourne, Parkville, Australia; i3S – Instituto de Investigação e Inovação em Saúde, Universidade do Porto, Porto, Portugal; CGPP-IBMC – Centro de Genética Preditiva e Preventiva, Instituto de Biologia Molecular e Celular, Universidade do Porto, Portugal; Division of Genetics, Oregon National Primate Research Center, Beaverton, USA; Genetics of Male Infertility Initiative (GEMINI) consortium; Institute of Reproductive Genetics, University of Münster, Münster, Germany; Centre of Reproductive Medicine and Andrology, University Hospital Münster, University of Münster, Münster, Germany; International Male Infertility Genomics Consortium (IMIGC); Department of Surgery (Urology), University of Utah, Salt Lake City, Utah, USA; Centre for Reproductive Health, Hudson Institute of Medical Research, Monash University, Clayton, Australia

**Keywords:** Dynein, protein transport, male infertility genetics, spermatogonia

## Abstract

Dynein complexes are large, multi-unit assemblies involved in many biological processes including male fertility via their critical roles in protein transport and axoneme motility. Previously we identified a pathogenic variant in the dynein gene *AXDND1* in an infertile man. Subsequently we identified an additional four potentially compound heterozygous variants of unknown significance in *AXDND1* in two additional infertile men. We thus tested the role of AXDND1 in mammalian male fertility by generating a knockout mouse model. *Axdnd1^-/-^* males were sterile at all ages but could undergo one round of histologically complete spermatogenesis. Subsequently, a progressive imbalance of spermatogonial commitment to spermatogenesis over self-renewal occurred, ultimately leading to catastrophic germ cell loss, loss of blood-testis barrier patency and immune cell infiltration. Sperm produced during the first wave of spermatogenesis were immotile due to abnormal axoneme structure, including the presence of ectopic vesicles and abnormalities in outer dense fibres and microtubule doublet structures. Sperm output was additionally compromised by a severe spermiation defect and abnormal sperm individualisation. Collectively, our data highlight the essential roles of AXDND1 as a regulator of spermatogonial commitment to spermatogenesis and during the processes of spermiogenesis where it is essential for sperm tail development, release and motility.

## Introduction

Sperm assembly begins in spermiogenesis, during which round spermatids are transformed into highly polarised cells, containing minimal cytoplasm, with the potential for motility and fertility (1). Given that nuclear gene transcription and translation ceases halfway through this process, the extreme metamorphosis of spermatids is achieved in large part through a series of coordinated protein and organelle transport pathways (2, 3). Multiple related transport pathways exist within haploid male germ cells, including along cytoplasmic microtubules through classical protein transport pathways, along the microtubules of the manchette via a process called intra-manchette transport (IMT) and along the microtubules of the axoneme via the classical, in addition to at least one modified, version of the intra-flagellar transport (IFT) pathway (2, 4). A family of complexes essential for many of these processes is the dyneins (5, 6). Dynein complexes are large multi-unit assemblies that play critical roles in protein transport and cell motility (7). There are two types of dynein complexes, the cytoplasmic dyneins, which transport cellular cargoes along microtubules towards their minus end within the cell cytoplasm (7–10), and axonemal dyneins, which drive the ATP-dependent beating of all motile cilia, including the sperm tail (11–13).

The canonical <u>cytoplasmic</u> dynein complex is composed of a dimer of two dynein heavy chains to which two intermediate chain, two light intermediate chain and three light chain dynein subunits binds (7, 14). In contrast to <u>cytoplasmic</u> dyneins, three heavy chain subunits define <u>axonemal</u> dynein complexes (α-, β- and γ-), alongside a unique set of light chain subunits not previously associated with the cytoplasmic dynein complex (15). In both classes of dynein complexes, the heavy chain dynein components act as the motor/engine of the complex. The heavy chains each possess a microtubule binding domain adjacent to an AAA module (15, 16), the latter which facilitates ATP hydrolysis to power transport along microtubules or cell motility. Following assembly of the core dynein complex, other regulatory proteins are recruited to enhance activity and define which cargoes (passengers [cytoplasmic]) or adapters bind to a specific dynein complex (17, 18). Different adapter (auxiliary) protein combinations exist within different tissues and are likely involved in different biological processes (19).

Previously we used whole exome sequencing on a group of infertile men suffering from non-obstructive azoospermia and identified a novel biallelic stop-gain variant in *AXDND1* (20). While AXDND1 contains a dynein light chain domain and is thus annotated as an axonemal dynein light chain-containing protein, it is notably different from the typical light chain dyneins. Specifically, it is considerably larger than *bona fide* mammalian dynein light chain proteins (∼8-30 kDa), at 120-125 kDa and does not encode other domains typical of light chain dyneins (8). AXDND1 does however contain a conserved PF10211 domain, homologous to the core dynein domains in DNALI1 and *Chlamydomonas* p28 (8, 21). While untested, we predict this domain likely confers AXDND1 the ability to interact with dynein heavy chains, the large motor protein subunits that drive cell motility or protein transport complex motility along microtubules (7). Previously it has been shown that the loss of *Axdnd1* function in mice results in male infertility due to its role in sperm head shaping via the manchette and tail assembly during spermiogenesis (22, 23), suggesting it is not exclusively a core axonemal dynein. This phenotype is a poor match with the azoospermic clinical presentation of the infertile man we identified to be carrying a stop-gain variant. To further test the role of AXDND1 in male fertility we therefore generated an *Axdnd1* knockout mouse line and robustly defined the effects of AXDND1 loss on germ cell development and male fertility.

In addition to a role in sperm tail development defined previously (23), we identify that AXDND1 is required for multiple aspects of male fertility, ultimately resulting in the loss of all germ cell types analogous to that observed in the full loss-of-function patient. Specifically, *Axdnd1^-/-^* males were sterile at all ages but were capable of undergoing one round of histologically complete spermatogenesis. The loss of AXDND1 resulted in a progressive loss of germ cell content and a breakdown of the blood-testis barrier that led to immune infiltration. Sperm produced during the first wave were immotile due to abnormal axoneme structure, including the presence of ectopic vesicles, loss of a portion of the outer dense fibres and microtubule doublets, abnormal mitochondrial sheath and sperm head formation, and poor sperm individualisation. Our data may suggest that AXDND1 functions in protein transport as an adapter in cytoplasmic dynein complexes, including in retrograde IFT during supplementation of the sperm tail axoneme with accessory structures, and in the processes of spermatid individualisation. Finally, we identified an additional four potentially compound heterozygous biallelic variants of unknown significance in *AXDND1*, in men suffering non-obstructive azoospermia.

## Results

### A loss of function variant in AXDND1 is associated with azoospermia and human male infertility

As part of the GEMINI study to define genetic causes of male infertility, we undertook exome sequencing of infertile men presenting with azoospermia or severe oligozoospermia and identified an azoospermic Portuguese man (patient 1) carrying a homozygous stop-gain variant in the dynein gene *AXDND1* (c.937T>C; p.R313X) (Figure 1A) (first reported in Nagirnaja et al. 2022). This variant affects exon 10 of 26 in *AXDND1*, a region that encodes a predicted axonemal dynein light chain (PF10211) domain. As identified by the variant effect predictor in gnomAD, the p.R313X variant is high confidence for a loss of AXDND1 function. The CADD score for this variant is 42, which places it in the top 0.01 % of deleterious variants in the genome. The predicted effect of this stop-gain mutation is nonsense-mediated decay of all the full-length RNA transcripts (*AXDND1*-201, 202 and 214; Supplementary Figure 1), resulting in a loss of *AXDND1* expression, i.e., it is equivalent to a full loss of function mutant in a mouse.

**Figure 1.**
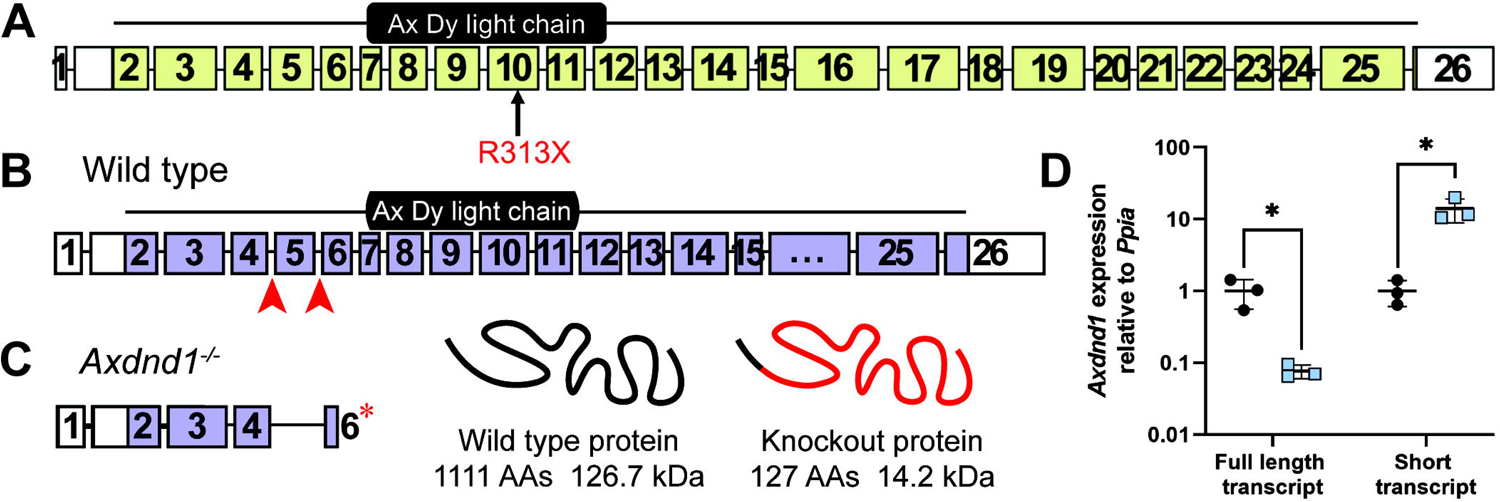
The *AXDND1* genetic variant and *Axdnd1*^-/-^ mouse model. **A.** We identified a high confidence disease-causing p.R313X stop-gain variant (red) in *AXDND1* in an infertile man with azoospermia. Human AXDND1 is encoded by 26 exons, with the stop-gain variant affecting exon 10, within the predicted axonemal dynein light chain domain. White exons are untranslated regions, green shaded exons are protein-coding. **B.** Full length mouse AXDND1 protein is encoded across 25 exons, with the dynein light chain domain encoded in exons 7-11. Red arrows denote guide RNA sites for generation of the knockout model via CRISPR. Purple shaded boxes represent protein-coding exons, while the white portion of exon 25 is the 3’ untranslated region. **C.** AXDND1 protein length in *Axdnd1^-/-^* mice, comprising of exon 2-4 and a truncated exon 6 with a premature stop codon. Wild type AXDND1 is represented in a basic form at 1111 amino acids in length and encoding 126 kDa of protein, while the red portion of the knockout protein represents the premature stop codon and the resulting 1064 amino acids that are not translated. **D.** Expression of the full length *Axdnd1* transcript and a smaller transcript not affected by the genetic modification were measured by qPCR, normalised to *Ppia* expression, * *p* < 0.05.

A comparison of the *AXDND1/Axdnd1* protein coding transcripts between human and mouse (Supplementary Figure 1) revealed that three transcripts in human (*AXDND1-*201, −2 and - 214) and four in mouse (*Axdnd1-*202, −2, −2 and −2) contained the PF10211 axonemal dynein light chain domain (exons 7-11). Despite encoding this predicted functional domain, transcript 214 in humans and 208 in mice encode truncated versions of the full length AXDND1 protein.

### AXDND1 is enriched in haploid male germ cells

We investigated *AXDND1* expression using published testis single cell sequencing data (24, 25). *AXDND1*/*Axdnd1* was expressed in all germ cell types, including spermatogonia, but most highly expressed within late spermatocytes and round spermatids in both men and mice (Figures 2A, B). These data are in agreement our qPCR expression analysis, wherein we observed an up-regulation of *Axdnd1* from postnatal day 14 to 20, corresponding to the first appearance of pachytene spermatocytes through to round spermatids. Assessment of *Axdnd1* expression across several mouse tissues revealed a clear enrichment in testes (Figure 2D), and an absence in other tissues investigated.

**Figure 2.**
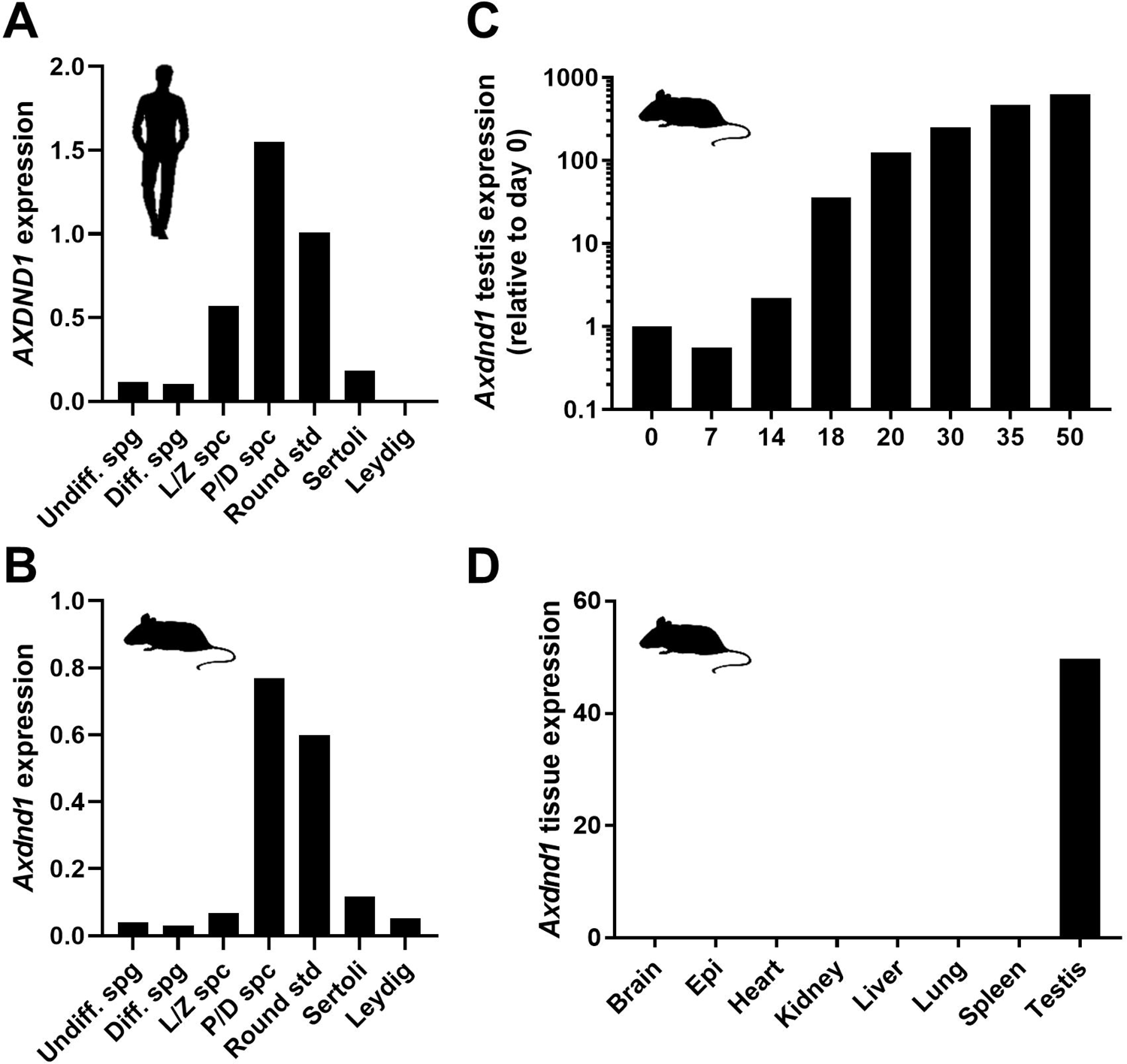
*AXDND1/Axdnd1* expression in testes and other organs. **A.** Human *AXDND1* and **B.** Mouse *Axdnd1* relative expression in testicular cells as identified by single cell RNA sequencing. L-R: undifferentiated spermatogonia, differentiated spermatogonia, leptotene and zygotene spermatocytes, pachytene and diplotene spermatocytes, round spermatids, Sertoli cells and Leydig cells. **C.** *Axdnd1* expression in mouse testes during the first wave of spermatogenesis (qPCR), normalised to *Ppia* expression and relative to day 0. **D.** *Axdnd1* expression across several mouse organs (qPCR), normalised to *Ppia* expression. Epi = epididymis.

### AXDND1 is essential for male fertility in mice

To test the role of AXDND1 in male fertility, we generated a knockout mouse model to recapitulate the predicted complete loss of function of the p.R313X *AXDND1* variant identified in infertile patient 1 (Table 1). We used CRISPR to remove exon 3 of the principal *Axdnd1* transcript and introduce a premature stop codon in exon 4 (Figure 1). Two founder mouse lines were generated. Following the confirmation of comparable phenotypes in both lines, our analyses focused on the *Axdnd1^del1^*line (described in the methods). Loss of expression of the full length *Axdnd1* transcript in this knockout line was confirmed via qPCR (Figure 1D), while the expression of a shorter transcript, encoded by exons 7-17 (*Axdnd1* transcript 208, Supplementary Figure 1) was upregulated approximately 10-fold (*p* < 0.05) compared to expression in wild type testes. Transcript 208 encodes a truncated protein of 55 kDa (compared with 126 kDa for the full-length protein) that contains the predicted PF10211 domain but is expected to be non-functional.

**Table 1.** Variants identified in AXDND1 in infertile men with azoospermia or severe oligozoospermia. Hom = homozygous, Het = heterozygous/phase unknown, LP = likely pathogenic, VUS = variant of unknown significance.

| Patient | Country of origin | Semen parameters and histology | AXDND1 variant |  |  | Pathogenicity |
| --- | --- | --- | --- | --- | --- | --- |
| 1 | Portugal | Non-obstructive azoospermia, no biopsy | c.937C>T | p.Arg313Ter | Hom | LP |
| 2 | Croatia | Non-obstructive | c.1607T>A | p.Leu536Gln | Het | VUS |
| M1557 |  | azoospermia, Sertoli cell-only | c.2602G>A | p.Ala868Thr | Het | VUS |
| 3<br>M2628 | Turkey | Non-obstructive azoospermia, elongating spermatids in < 50% tubules | c.2451A>C | p.Lys817Asn | Het | VUS |
|  |  |  | c.2843A>G | p.Asp948Gly | Het | VUS |

Fertility of *Axdnd1^-/-^* males was tested at two ages: 1) at the completion of the first wave of spermatogenesis and epididymal maturation (42-50 days of age) and 2) after several rounds of spermatogenesis (70-80 days of age). At both ages, *Axdnd1*^-/-^ males were sterile (Figure 3A; *p* < 0.0001) although mating appeared normal as evidenced by the presence of copulatory plugs (0 pups/plug). Wild type males plugged and sired several litters (average of 8.25 ± 0.5 and 7.36 ± 1.1 pups/plug, at postnatal days 42-50 and 70-80 respectively) at both ages. No differences were found in the body weight of *Axdnd1*^-/-^ males compared to wild type across all ages investigated (Figure 3B).

**Figure 3.**
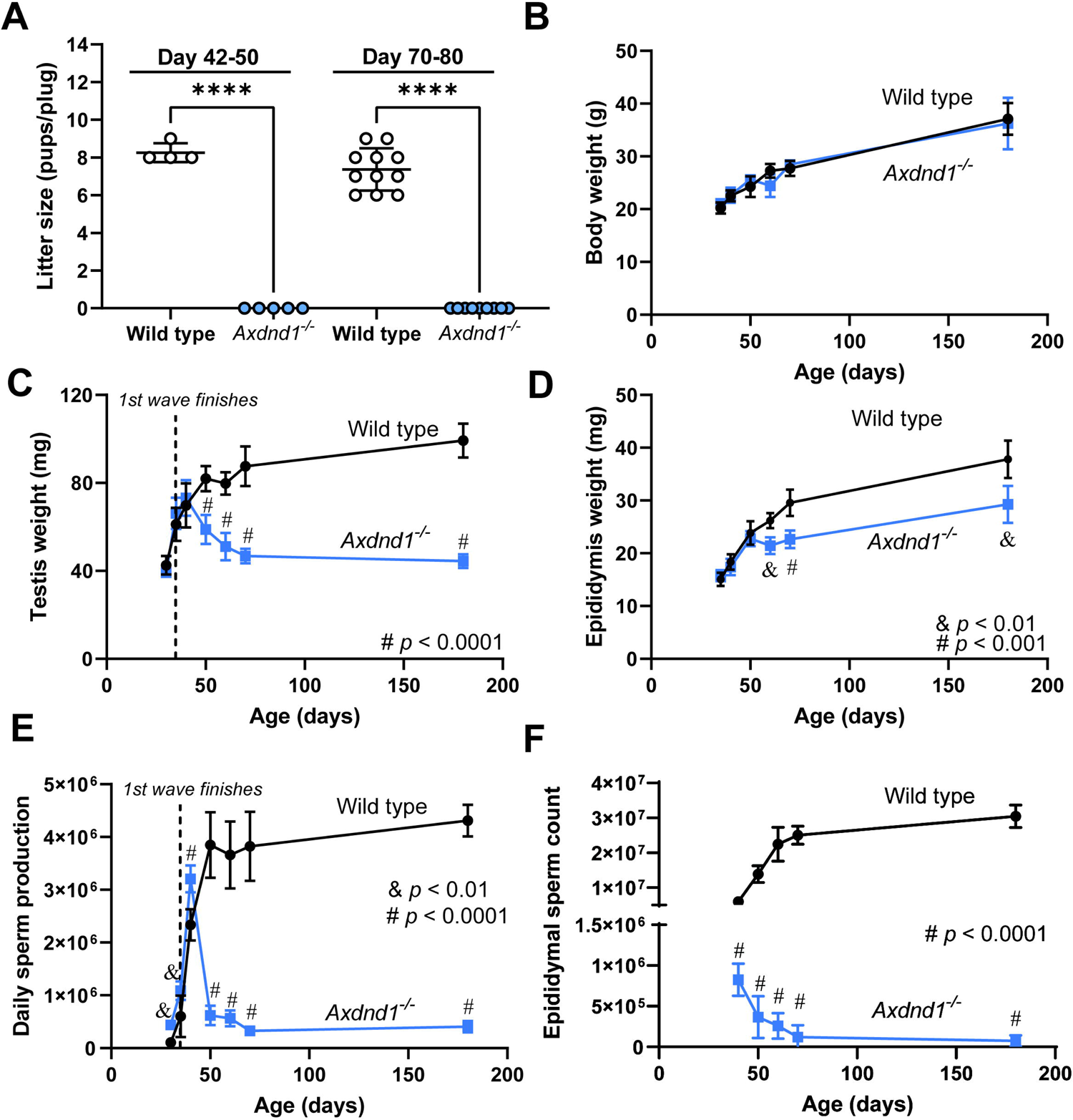
AXDND1 is essential for male fertility and normal sperm production capacity. Male reproductive parameters and body weight of wild type and *Axdnd1*^-/-^ males were assessed across an age range of 35-180 days of age **A.** Male fertility as assessed by litter size for males mated just after the first wave of spermatogenesis (42-50 days of age) or after multiple waves. **B.** Body weight. **C.** Testis weight, including day 30 data points **D.** Epididymis weight. **E.** Daily sperm production, include day 30 data points. **F.** Epididymal sperm count. & = *p* < 0.01; # = *p* < 0.0001 at respective age point.

Next, reproductive parameters of wild type and *Axdnd1*^-/-^ males were assessed over an age range of 40-180 days of age (Figure 3) to explore the role of AXDND1 in establishing the first wave of spermatogenesis and in the maintenance of spermatogenesis. Testis weights were comparable between wild type and *Axdnd1*^-/-^ males until 40 days of age, after which a significant divergence was observed (Figure 3C; *p* < 0.0001). *Axdnd1*^-/-^ testes weights progressively declined until 70 days of age, then remained constant through to 180 days of age. At postnatal day 180, *Axdnd1*^-/-^ testis weight was 45% of that measured from wild type males. In accordance, epididymis weights were comparable between genotypes up until day 50 of age, then diverged from day 60 of age (Figure 3D; *p* < 0.01). Such changes were suggestive of the successful completion of one round of spermatogenesis followed by an age-related degeneration. To explore this possibility quantitatively, sperm production was measured as a function of age (Figure 3E). During the first wave of spermatogenesis (30 and 35 days of age), knockout males generated significantly more sperm compared to wild type (*p* < 0.01). At 40 days of age, knockout males produced 30% more sperm than age matched wild type males (*p* < 0.0001). By 50 days of age, however, *Axdnd1*^-/-^ daily sperm production, decreased to 19% of wild type levels (*p* < 0.0001). Similar levels of sperm production were maintained through to 180 days of age. Despite no initial increase in sperm number, an analogous pattern of reduced sperm number was observed within the epididymis as a function of age. At day 40 of age, sperm counts in *Axdnd1*^-/-^ epididymides were 12% of wild type levels (*p* < 0.0001). Epididymal sperm counts then declined with age, and at 180 days of age were only 0.33% of wild type levels. The disconnect in daily sperm production in the testis and epididymal sperm at day 40 is suggestive of a widespread spermiation defect. In support of this hypothesis, significant numbers of sperm were observed within stage IX-X tubules of day 40 *Axdnd1*^-/-^ testes (Supplementary Figure 2B).

A comparable phenotype was observed in the second *Axdnd1*^-/-^ line (*Axdnd1^del2^*, Supplementary Figure 3), thus confirming that the phenotype was specific to the loss of AXDND1 function and not an undetected ‘off target’ CRISPR-mediated loss of gene function somewhere else in the genome.

To explore the consequences of AXDND1 loss at a cellular level, we investigated testis histology (Figure 4) in wild type and *Axdnd1*^-/-^ males at 40 (just following the first wave of spermatogenesis), 70 (after several waves) and 180 (aged males) days of age. Analogous to the situation in wild type mice (Figure 4A), all adult germ cell types were present at 40 days of age in *Axdnd1*^-/-^ testes (Figure 4B). By 70 days of age, however, and consistent with the changes in testis weight, spermatogenesis was overtly compromised as highlighted by a significant reduction in, or complete absence of, germ cells in a portion of seminiferous tubules, resulting in vacuoles in the seminiferous epithelium (Figure 4D, red arrowheads). Associated with this was an influx of cells to the interstitial compartment of the testis (Figure 2D, red dashed lines), which were confirmed to be immune cells by staining for CD45 (Supplementary Figure 2D). These disruptions to spermatogenesis in knockout testes worsened by 180 days of age (Figure 4F), as characterised by a significant increase in the incidence of Sertoli cell-only tubules (Figure 4J, *p* < 0.0001).

**Figure 4.**
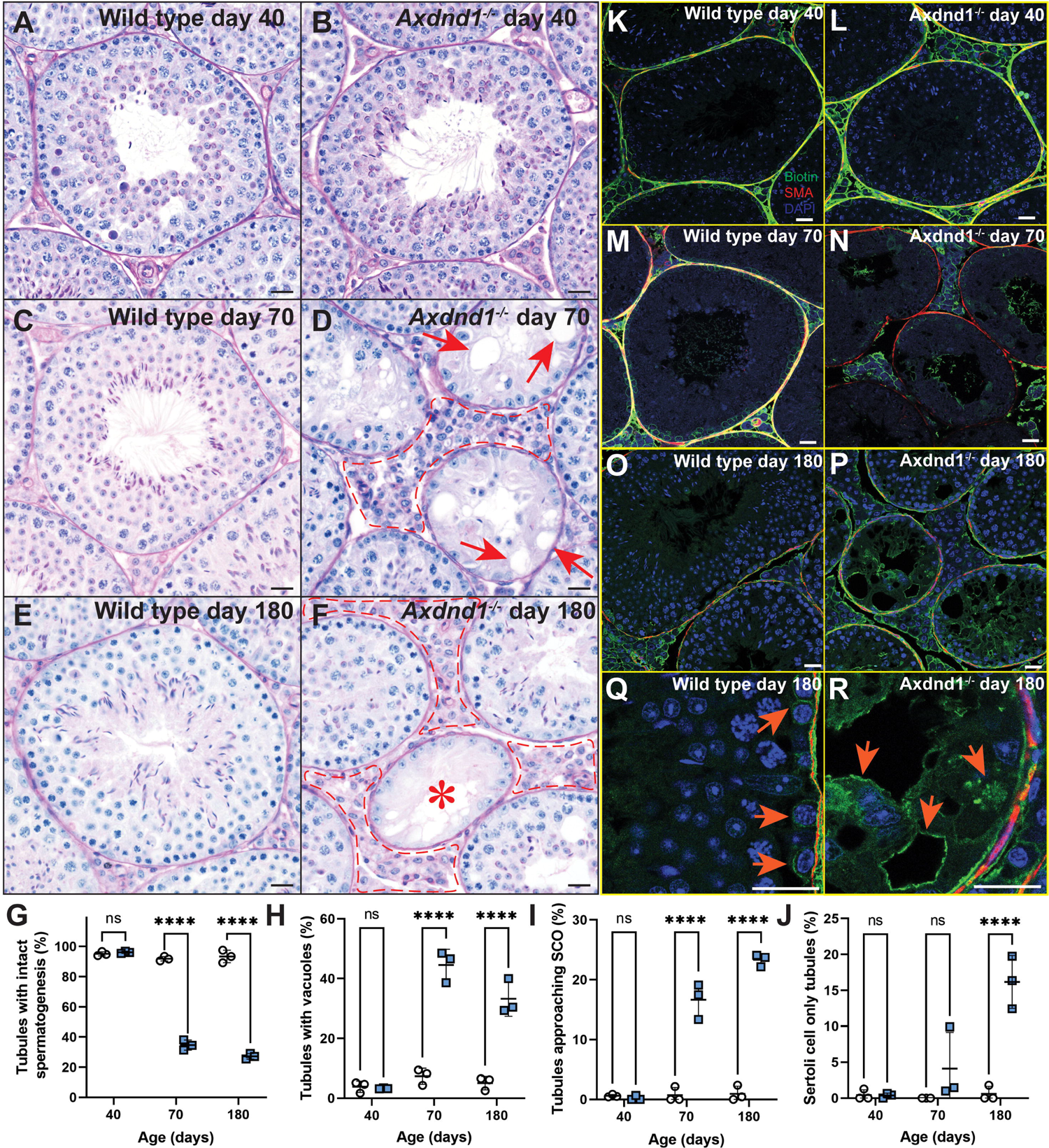
AXDND1 is required for normal and sustained spermatogenesis. Testis histology and apoptosis was assessed at days 40, 70 and 180 to investigate the role of AXDND1 in spermatogenesis. **A.** Wild type and **B.** *Axdnd1^-/-^* knockout testes at day 40 of age. **C.** Wild type and **D.** *Axdnd1^-/-^* knockout testes at day 70 of age. Arrows point to vacuoles in the seminiferous epithelium. Red dashed lines denote abnormal interstitium. **E.** Wild type and **F.** *Axdnd1^-/-^* knockout testes at day 180 of age. The asterisk denotes a Sertoli cell-only tubule, devoid of germ cells. Scale bars are 20 µm in length for all panels. **G.** Assessment of seminiferous tubules with no abnormal presentation (intactness). **H.** Assessment of tubules with vacuoles **I.** Assessment of tubules with severe germ cell loss, approaching a Sertoli cell-only phenotype **J.** Assessment of tubules with a Sertoli-cell only phenotype. **K.** Assessment of apoptosis via caspase staining. * = *p* < 0.05, **** = *p* < 0.0001.

To quantify these changes in spermatogenic quality, the proportion of tubules presenting with complete spermatogenesis, hypospermatogenesis, or a Sertoli cell-only epithelium was quantitated (Figure 4G-J). These data revealed that the proportion of normal tubules with intact spermatogenesis decreased between 40 and 70 days of age in *Axdnd1^-/-^* testes (Figure 4G, *p* < 0.0001) and accordingly the proportion of tubules with vacuoles and tubules approaching Sertoli cell-only (severe hypospermatogenesis) in *Axdnd1*^-/-^ testes were both significantly elevated from 70 days of age (Figures 4H, I; *p* < 0.0001). As a measure of complete germ cell loss, the number of complete Sertoli cell-only tubules was significantly elevated at 180 days of age (Figure 4J, *p* < 0.0001). In agreement with this reduction in germ cell content, the quantitation of germ cell apoptosis via caspase staining revealed a significant increase in apoptotic germ cells in *Axdnd1^-/-^* testes at 70 days of age compared to wild type (*p* = 0.02; Supplementary Figure 2E).

As described in more detail below, those sperm present within the epididymis were immotile, thus explaining the origin of infertility in young adult males (Figure 6). In addition to the severe tail defects, the loss of AXDND1 resulted in a significant increase in head shape abnormality (Supplementary Figure 4). Objective sperm head shaping analysis revealed a reduction in the percentage of sperm heads classified as most normal (cluster 1, *p* = 0.0033) and an increase in the abnormal clustering (cluster 3, *p* = 0.023). This finding is in support of a critical role of AXDND1 in manchette dynamics (22, 23) and its localisation to the manchette.

### The blood-testis barrier is compromised in Axdnd1^-/-^ testes

The accumulation of immune cells within the interstitial space in older *Axdnd1^-/-^* (Supplementary Figure 2C) males suggested the loss of patency of the blood-testis barrier. The blood-testis barrier is a series of intercellular junctions between adjacent Sertoli cells that functions to protect the immunologically foreign meiotic and haploid germ cells from immune system attack (26). To directly test the patency of the blood-testis barrier we incubated testes with a biotin tracer (Figure 4K-R). This approach revealed that the blood-testis barrier was intact in *Axdnd1*^-/-^ testes at 40 days of age as evidenced by the localisation of biotin exclusively within the interstitium and the basal compartment of the seminiferous epithelium which is external to the blood-testis barrier. By contrast, in 70- and 180-day old *Axdnd1*^-/-^ testes the biotin tracer permeated deep within the tubules (Figure 4N, P, R). As expected, the blood-testis barrier was intact in wild type testes at all ages (Figure 4M, O, Q). These data reveal that the loss of AXDND1 leads to a loss of blood-testis barrier patency. It is unclear if this is a primary or secondary consequence of AXDND1 loss. Regardless, the loss of blood-testis barrier patency would be expected to accelerate the seminiferous tubule degeneration with increasing age.

### AXDND1 is required for normal sperm tail development and function

The presence of sperm within day 40 *Axdnd1^-/-^* testes and epididymides suggested the possibility of fertility in young animals. However, as described above, the mating experiments revealed males were sterile. To define the origin of this infertility, epididymis histology was examined at 40 days of age (Figure 5). As outlined above, while sperm were present in the epididymis, numbers were decreased compared to age-matched wild type males. By 70 days of age (Figure 5D, H) very few sperm were seen and, instead, the cauda contained many round cells – as evidenced by both prematurely sloughed germ cells as confirmed previously (23) and likely immune cells (Supplementary Figure 2G). This finding was consistent in aged males at 180 days of age (Figure 5F), where widespread phagocytosis of spermatid nuclei by immune cells was evident.

**Figure 5.**
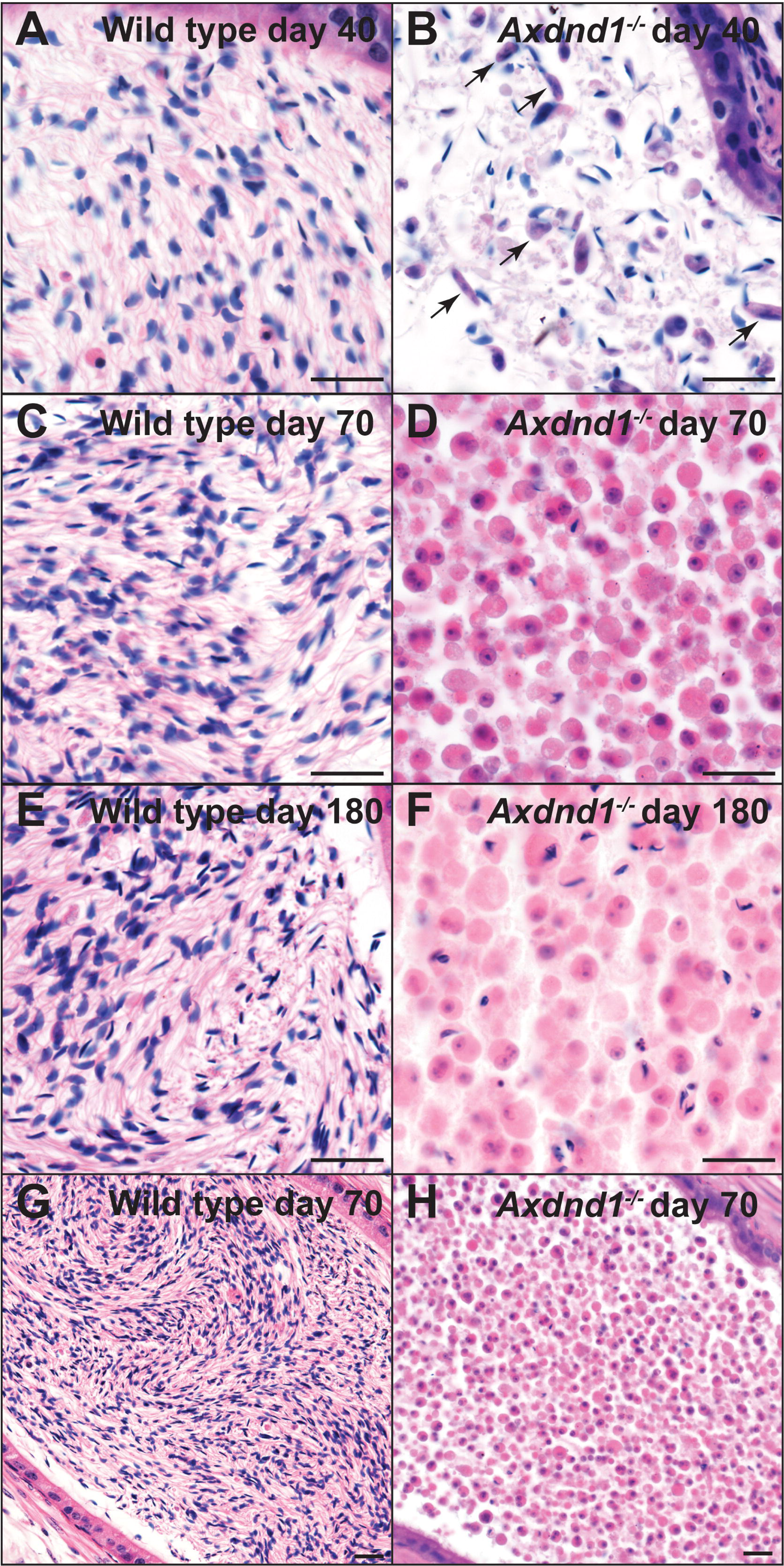
AXDND1 is required for normal sperm release into the epididymis. Cauda epididymis histology was assessed at days 40, 70 and 180 of age to investigate the cell types present. **A.** Wild type and **B.** *Axdnd1^-/-^* knockout epididymides at day 40 of age. **C.** Wild type and **D.** *Axdnd1^-/-^*knockout epididymides at day 70 of age. **E.** Wild type and **F.** *Axdnd1^-/-^* knockout epididymides at day 180 of age. Lower power images of **G.** Wild type and **H.** *Axdnd1^-/-^* knockout epididymides at day 70 of age are also shown. Scale bars are 10 µm in length.

Of those sperm released by *Axdnd1*^-/-^ testes at day 40, the majority were abnormal, as highlighted by highly coiled sperm tails (Figure 5B, arrows; Figures 6A, F). Most possessed an abnormal midpiece region, and/or sperm that were not individualised correctly and signs of abnormal head shaping. Sperm isolated from the cauda epididymis of *Axdnd1^-/-^* males were completely immotile (Figure 6B, *p* < 0.0001). Further, motility was not recovered with the addition of membrane permeable ATP, suggesting a core structural defect within the axoneme (2). An analysis of midpiece / mitochondrial defects revealed that the mitochondrial sheath of sperm was poorly formed or missing in 87% of sperm from *Axdnd1^-/-^* males but only 5% of wild type sperm (*p* < 0.0001; Figure 6C). Sperm tail length was measured in individual sperm, noting this was not possible for those with coiled tails, and revealed no significant differences between genotypes (Figure 6D).

**Figure 6.**
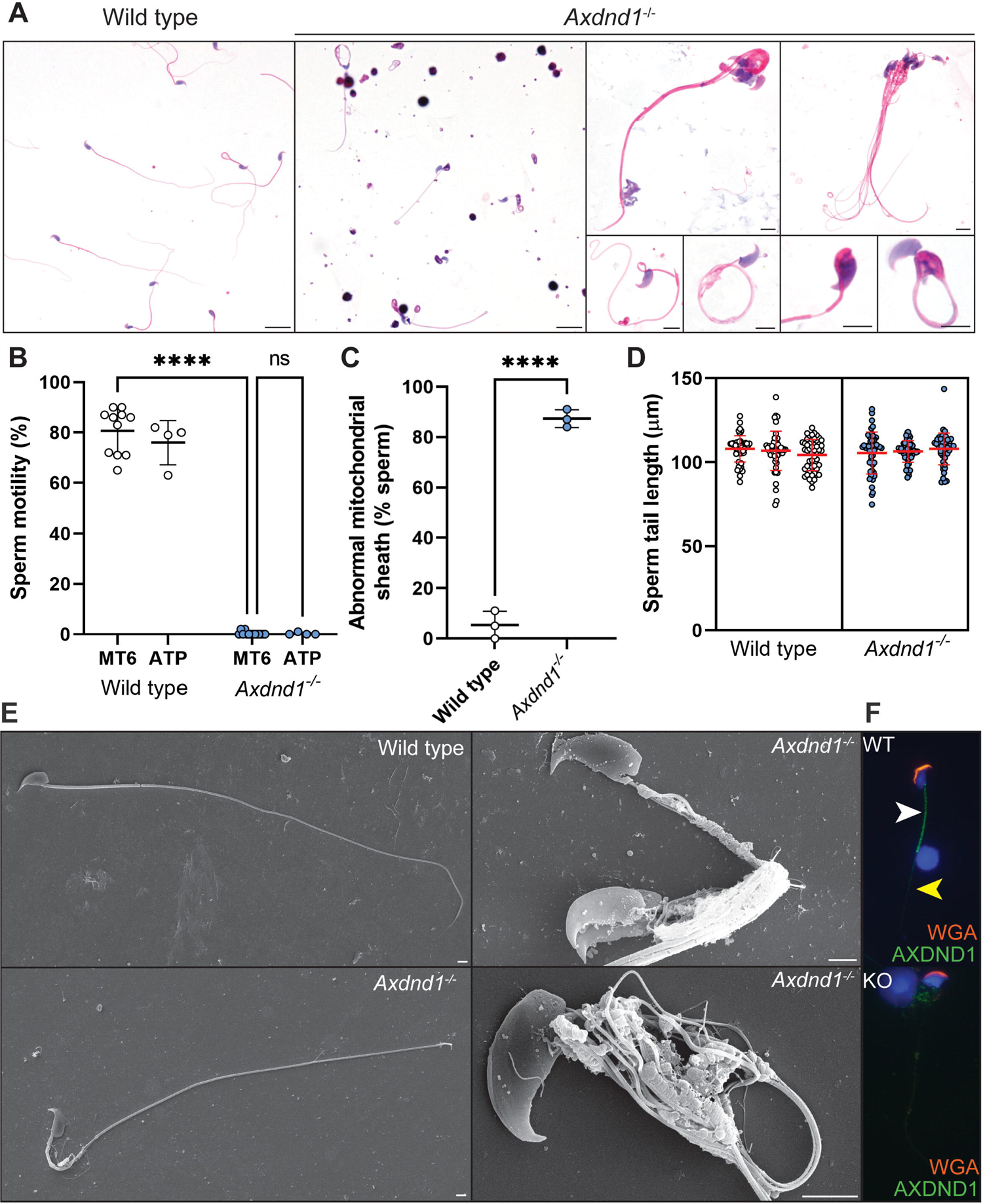
AXDND1 is an essential regulator of sperm tail development and required for sperm motility. **A.** Sperm morphology of sperm from wild type (left panel) and *Axdnd1^-/-^* (all other panels) stained with haematoxylin and eosin. Scale bars in large panels are 10 µm in length and in small panels are 5 µm in length. **B.** Sperm motility in basal medium and with the presence of exogenous ATP. **C.** Quantification of sperm mitochondrial sheath abnormalities. **D.** Sperm tail length. **E.** Scanning electron microscopy of sperm. Lower power panels (left) depict whole sperm external structure (plasma membrane removed). Higher power panels (right) focus on sperm midpiece / mitochondrial sheath regions in knockout. Scale bars are 2 µm in length. AXDND1 localisation in sperm collected via squash preparations (green), counterstained with DAPI (blue) and wheat germ agglutinin (acrosome marker, red). A white arrow denotes midpiece staining and a yellow arrow points to punctate principal piece staining. A negative control (sperm from knockout) is shown in the lower panel with no AXDND1 staining. **F.** AXDND1 localisation in sperm collected via squash preparations (green), counterstained with DAPI (blue) and wheat germ agglutinin (acrosome marker, red). A white arrow denotes midpiece staining and a yellow arrow points to punctate principal piece staining. A negative control (sperm from knockout) is shown in the lower panel with no AXDND1 staining.

Examination of cell ultrastructure via scanning electron microscopy on membrane stripped sperm revealed that sperm generated by *Axdnd1^-/-^* males had highly abnormal mitochondrial sheath structure and mitochondrial morphology (Figure 6E). In contrast to the neatly coiled mitochondria of sperm from wild type mice, those from *Axdnd1^-/-^* males, possessed poorly coiled mitochondria and gaps within the sheath where no mitochondria were present. In the midpiece compartment, microtubules and outer dense fibres splayed outward from the axoneme core and in some cases appeared to have snapped off from the tail. As indicated above, sperm were poorly individualised, highlighting a failure of cytoplasm removal (part of the spermiation process) or a failure to remove intracellular junctions between individual spermatids during the later steps of spermiogenesis. By contrast, the structure of the fibrous sheath of these sperm appeared to be largely normal as observed externally from a scanning electron microscopy level, but also internally as investigated by transmission electron micrographs (Figure 7E, H, I).

**Figure 7.**
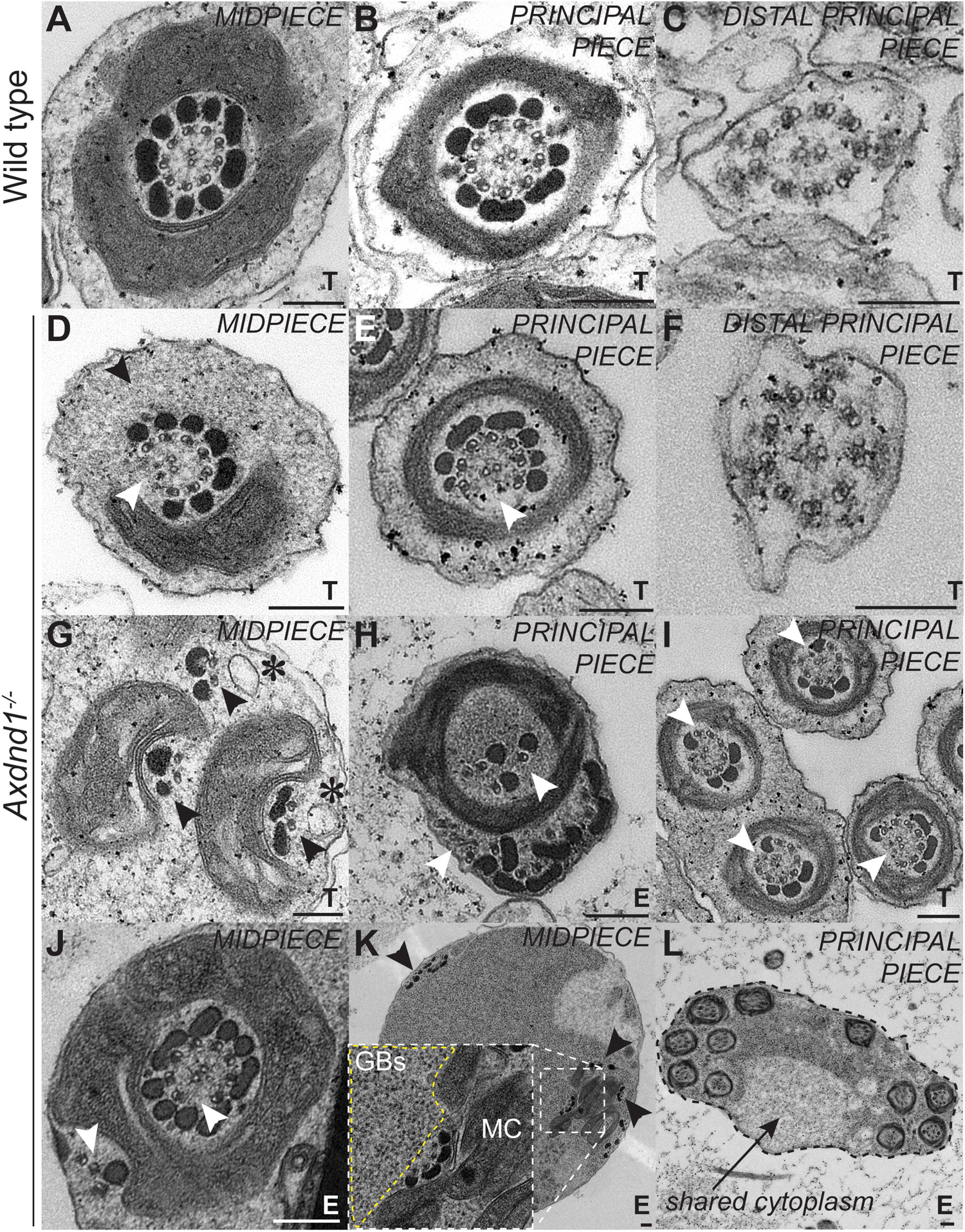
AXDND1 is a critical regulator of sperm axoneme ultrastructure. Transmission electron microscopy images of sperm axoneme cross-sections across all three regions of the sperm tail. Testis (T) versus epididymis (E) sperm samples are designated at the bottom right corner of each panel. **A-C.** Step 16 spermatids from wild type and **D-F.** *Axdnd1^-/-^* at the midpiece, principal piece and end piece level, respectively. The black arrow in D denotes missing mitochondria, and in D+E white arrows point to missing microtubule doublets and outer dense fibres (ODFs). **G-L.** Various defects in sperm tail ultrastructure in sperm from *Axdnd1^-/-^*. **G.** Black arrows points to separate groups of microtubules and ODFs associated with mitochondria; asterisks denote vesicles associated with these microtubules. **H.** White arrows point to disordered microtubules, including those displaced from the core exoneme encased within the fibrous sheath. **I.** White arrows point to missing microtubules and ODFs. **J.** White arrows point to displaced microtubules. **K.** Black arrows point to disordered microtubules and ODFs throughout the cytoplasm. An insert shows granulated bodies (GBs, yellow dashed line) accumulated next to four microtubule doublets and ODFs associating with mitochondria (MC). **L.** A black dashed line and arrow denotes a shared cytoplasm among several individual sperm tails. Scale bars are 200 nm in length.

To define why sperm from *Axdnd1^-/-^* males were immotile, we used transmission electron microscopy on developing spermatids (testis) and released sperm (epididymal sperm) (Figure 7). An investigation of step 15-16 spermatids, just before sperm release, and epididymal sperm from *Axdnd1*^-/-^ males revealed a battery of axoneme and other sperm tail defects (Figure 7D-F) compared to developmentally matched cells from wild type males (Figure 7A-C). In most sperm, a portion of axoneme microtubule doublets were missing or ectopically positioned in both the midpiece and the principal piece. In addition, abnormal ODF loading was observed in virtually all cross-sections (Figures 7D-E, G-J), as highlighted by absent or small ODF structures. Further, ectopic vesicles were identified in the axoneme (Figure 7G, asterisks) and as outlined above, abnormal mitochondrial loading (Figure 7D, G) was observed in the midpiece. Collectively, these defects raise the possibility of a defect in protein and/or organelle transport into the developing sperm tail compartment. In support of such defects, we observed the over-accumulation of granulated bodies (precursor ODFs) in the cytoplasm (Figure 7K insert – yellow dashed line; Figure 7L). As evidenced in distal principal piece sections, core axoneme ultrastructure appeared broadly normal in sperm from *Axdnd1^-/-^* males (Figure 7F), suggesting defects were exacerbated during the loading of accessory structures. Consistent with the finding of failed sperm individualisation, we detected many examples of multiple sperm sharing a cytoplasm in samples from the epididymis (Figure 7L).

To further explore if AXDND1 might play a direct role in organelle/protein transport within the sperm tail, we stained spermatids from wild type and knockout males with an AXDND1 antibody. While the antibodies we tested did not result in specific staining in formaldehyde fixed testis sections they did appear to be specific in elongating spermatids harvested using the dry down method and fixed with paraformaldehyde (27). As shown in Figure 6F, AXDND1 localised strongly to the midpiece region and in a punctate manner in the principal piece in developing wild type spermatids but not in knockout spermatids. AXDND1 also localised to the manchette and the developing head-tail coupling apparatus of spermatids (Supplementary Figure 2F).

### AXDND1 is a regulator of spermatogonial commitment to spermatogenesis

The elevated daily sperm production measured in *Axdnd1*^-/-^ testes at days 30, 35 and 40 of age followed by a rapid decline is suggestive of AXDND1 playing a role in maintaining the balance between spermatogonial stem cell self-renewal and commitment to spermatogenesis. To explore this, we quantified the number of round spermatid numbers at day 22 of age (Figure 8A), where spermatids first appear at postnatal day 20 in wild type males (28). This analysis revealed a significant increase in round spermatid number in seminiferous tubules in *Axdnd1^-/-^* testes compared to those from wild type animals (Figure 8B, *p* = 0.0026) and, accordingly, a significant increase in the average tubule area (Figure 8C, *p* = 0.0071).

**Figure 8.**
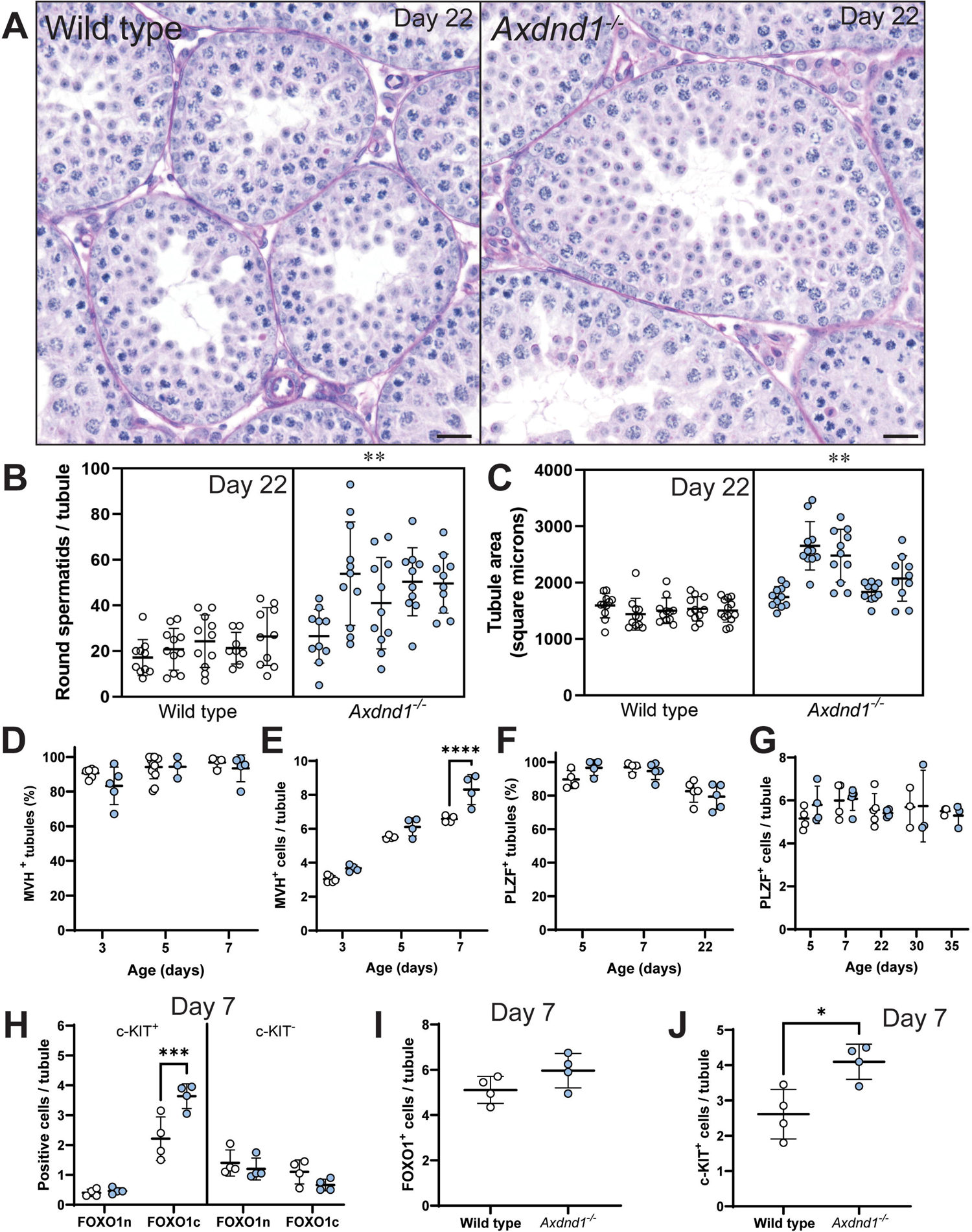
AXDND1 regulates spermatogonial commitment during the first wave of spermatogenesis. **A.** Testis histology at postnatal day 22 of age, depicting larger tubules and elevated round spermatid numbers in *Axdnd1^-/-^* testes. **B.** Round spermatid counts per tubule and **C.** Tubule area at postnatal day 22. **D.** MVH^+^ tubules and **E.** Average MVH^+^ cells per tubule at postnatal days 3, 5 and 7. **F.** PLZF^+^ tubules and **E.** Average PLZF ^+^ cells per tubule at postnatal days 3, 5 and 7. **H.** Average numbers of FOXO1 nuclear (n) and cytoplasmic (c) cells, separated into both c-KIT^+^ (committed) and c-KIT^-^ spermatogonia populations. **I.** Average FOXO1^+^ stained cells and **J.** c-KIT^+^ cells per tubule. * = *p* < 0.05, ** = *p* < 0.01, *** = *p* < 0.001, **** = *p* < 0.0001. Scale bars = 20 µm.

To define the origins of the increase in cell number, we quantified the numbers of MVH^+^ germ cells (pan germ cell marker) at postnatal day 3-7 and PLZF^+^ cells (undifferentiated and early differentiating spermatogonia) at postnatal day 5, at a time when nascent undifferentiated and differentiating spermatogonia have been generated from fetal germ cell precusors (pro-spermatogonia) (29), through to day 35 of age. Differentiating spermatogonia generated from pro-spermatogonia produce the first wave of spermatogenesis while subsequent sperm production is dependent on the undifferentiated pool that contains spermatogonial stem cells. At post-natal day 7 normal numbers of tubules were positive for MVH^+^ staining (Figure 8D), but the number of germ cells per tubule was increased in *Axdnd1*^-/-^ testes (Figure 8E, *p* < 0.0001). No differences were seen in the number of PLZF^+^ spermatogonia at all ages assessed (Figure 8F, G). To extend this analysis, we investigated staining for 1) c-KIT, a marker of undifferentiated spermatogonia, and 2) the transcription factor FOXO1, a marker of pro-spermatogonia and undifferentiated spermatogonia (30). FOXO1 is predominantly cytosolic in neonatal gonocytes, then translocates to the nucleus in undifferentiated spermatogonia but becomes cytosolic then downregulated upon spermatogonial differentiation (30, 31). We identified a significant increase in the number of c-KIT positive spermatogonia with a FOXO1 cytoplasmic localisation, i.e., spermatogonia at early differentiation stages (Figure 8H, *p* < 0.001) at day 7 of age in *Axdnd1^-/-^* testes. Total numbers of c-KIT positive spermatogonia were also increased (Figure 8J, *p* = 0.014). Consistent with unaltered numbers of PLZF^+^ spermatogonia, we did not identify a significant change to overall numbers of FOXO1^+^ germ cells *Axdnd1^-/-^* testes (Figure 8I, *p* = 0.13). Collectively, these data suggest that while AXDND1 is not required for spermatogonial development, it is a regulator of commitment of the spermatogonia originating from gonocytes to spermatogenesis during the first wave of spermatogenesis and that in its absence, an elevated number of nascent spermatogonia commit to differentiate to become sperm.

### Human male infertility clinical validity assessment for AXDND1

In order to search for additional men carrying AXDND1 variants, cohorts of men within the International Male Infertility Genomics Consortium (imigc.org) were assessed. We identified four variants (Table 1): two nonsynonymous SNVs in a Croatian man (patient 2, M1557) with azoospermia and a Sertoli cell-only presentation, and two nonsynonymous SNVs in a Turkish man with azoospermia (patient 3, M2628), where a biopsy revealed less than 50% of seminiferous tubules bearing elongating spermatids. All four of these variants were classified as variants of unknown significance. While the phases could not be determined in both cases (segregation analysis and long read sequencing), the disease presentation is consistent with patient 1 wherein a complete loss of *AXDND1* function variant was identified.

## Discussion

Herein our data confirm that AXDND1 is essential for male fertility for mice and men, functioning in several phases of spermatogenesis. AXDND1 plays a critical role in regulating the balance of spermatogonial commitment versus renewal during the first wave of spermatogenesis. It is then required for the formation of functional sperm tail structures, sperm individualisation and head shaping, and spermiation. These latter functions are consistent with a role in regulating protein transport along microtubule structures. To make things worse, while mice are still relatively young, the absence of AXDND1 leads to a loss of blood-testis barrier patency and the recruitment of immune cells into the testis and seminiferous epithelium further exacerbating pathology. While previous studies have identified AXDND1 to be essential for male fertility in mice, they failed to identify the evolving phenotype of *Axdnd1^-/-^* mice, which exacerbates with age (22, 23).

AXDND1 was originally annotated as a light chain-domain containing protein of the axonemal dynein class. If this was the case, it would be expected that AXDND1 plays a specific and essential role in axoneme function to manifest sperm motility. We did observe a complete loss of motility in the absence of AXDND1. We, however, also identified several structural defects in sperm from *Axdnd1^-/-^* males which suggest AXDND1 plays a more general role in protein or organelle transport. AXDND1 contains a PF10211 domain that is conserved in DNALI1, a dynein light intermediate chain which has been shown to interact with cytoplasmic dynein 1 (21). Therefore, AXDND1 may play dual roles in dynein complexes to accomplish axoneme-mediated cell motility and for cytoplasmic dynein-mediated transport. A similar hypothesis has been raised for the light intermediate dynein chain DNALI1 (32). A variant encoding hypomorphic DNALI1 in mice resulted in axoneme microtubule doublet and fibrous sheath abnormalities (32). Alternatively, the motility defect observed in sperm from *Axdnd1^-/-^* males may be exclusively due to the severe ultrastructural defects, including the partial loss of microtubule doublets and outer dense fibres throughout the axoneme. These data suggest AXDND1 may act as an adapter to optimise the interaction between dynein complexes and cargoes (e.g., ODFs) to allow their efficient transport along the microtubules of the manchette to the site of entry of the sperm tail compartment (cytoplasmic dynein 1) and/or via retrograde transport along the developing sperm tail (cytoplasmic dynein 2) in the ciliary compartment. The additional presence of abnormal concentrations of granulated bodies in the cytoplasm of *Axdnd1^-/-^* spermatids, indicative of poor entry to the site of the sperm tail, may also point to a role for AXDND1 in cytoplasmic dynein 1. AXDND1 does not appear to play a role in core intraflagellar transport that seeds the axoneme core of the sperm tail as identified by normal sperm tail lengths and a normal axoneme presentation in the end piece of sperm from the knockout model. The sperm tails were, however, immotile, and unresponsive to cell-permeable ATP, supporting defective assembly.

The loss of sperm individualisation has rarely been seen in mouse models of male infertility. An additional light chain domain containing dynein has, however, previously been shown to be essential for the process of sperm individualisation in flies (33), suggesting a potential conserved role for dyneins in this process. Recently, it was demonstrated that the process of sperm individualisation in mice is driven by an autophagy pathway (34) and likely requires efficient neddylation, a process of targeting cellular material for degradation that is similar to ubiquitination (35). Whether AXDND1 aids in trafficking members of either pathway to sites for targeted removal of the cytoplasmic bridges and other material required for individualisation should be tested.

Herein where we describe AXDND1 localises to the manchette, and in previous studies, it is clear that AXDND1 is required for normal manchette structure and thus sperm head shaping (22). Taken together these data point to a direct role for AXDND1 at the manchette. Of note, the manchette provides dual functions in head shaping and as a scaffold for intramanchette protein and vesicle transport (2), whereby cellular cargoes are trafficked from the Golgi or cytoplasm to a region near the entry to the sperm tail compartment. It is therefore possible that at least part of the ODF developmental defects may have arisen due to a reduction in intramanchette transport (36–38).

Although the mechanisms of mitochondrial loading into the sperm midpiece are poorly understood, it is hypothesised that the manchette also aids as a scaffold to align the mitochondria parallel to the axoneme prior to their loading onto the ODFs (2). The midpiece presentation in sperm from *Axdnd1*^-/-^ males is, however, more severe than that previously identified in knockout models wherein the manchette is dysfunctional (3, 39). Of note, the severe mitochondrial sheath defects in *Axdnd1*^-/-^ males bear resemblance to sperm from models where sperm individualisation is defective (34, 35), raising the possibility it is a secondary effect of failed individualisation. However, given that defects were evident during mitochondrial sheath development in spermatids, and the localisation of AXDND1 to the developing spermatid midpiece, it is more likely that AXDND1 plays a direct role in facilitating the loading of mitochondria into the midpiece compartment.

Finally, and importantly we demonstrated a previously unappreciated role for AXDND1 as a regulator of spermatogonial commitment to spermatogenesis. The overcommitment of spermatogonia in *Axdnd1*^-/-^ males resulted in an elevated sperm number during the first wave of spermatogenesis at the expense of subsequent waves. How AXDND1 represses the commitment of spermatogonia to spermatogenesis is unclear but may be explained by it playing a role in the transport of specific cell-regulatory factors. For example, previous research has shown that the RNA-binding protein DAZL, which is essential for the translation of RNAs important for male germ cell development, interacts directly with the cytoplasmic dynein complex (40). In addition, it was postulated that RNAs bound to DAZL could be transported by dyneins when required for developmental processes and protein translation (40). AXDND1 may be involved in a similar process, acting as a dynein adapter required for specific germ cell regulatory factors.

One caveat of this study was the inability to phase the potential compound heterozygous variants in *AXDND1* patients 2 and 3. In support of their potential pathogenicity, however, the clinical presentation of both men was azoospermia, which is a match with patient 1 and the *Axdnd1* knockout mouse. Patient 3 presented with severe hypospermatogenesis and patient 2 with Sertoli cell only tubules. Hypospermatogenesis leading to an eventual Sertoli cell only phenotype was observed in the knockout mouse model, highlighting the benefits of studying the evolving phenotype with age.

In this study we have established AXDND1 as a key regulator of spermatogonial commitment, sperm tail development and male fertility. We describe one high confidence genetic variant and four others of unknown significance in *AXDND1* in infertile males with azoospermia. Our data suggest AXDND1 likely functions in the cytoplasmic dynein complex in male germ cells, in contrast to its predicted role as an axonemal dynein.

## Materials and methods

### Mouse model production

The structure for human *AXDND1* and mouse *Axdnd1* transcript information (ENSG00000162779 and ENSMUSG00000026601, respectively) are shown in Supplementary Figure 1 and data was sourced from Ensembl. To test the requirement for *AXDND1* in male fertility, *Axdnd1* knockout (*Axdnd1^-/-^)* mice were generated by the Monash Genome Modification Platform (MGMP; Monash University, Australia) using CRISPR/Cas9 technology on C57BL/6J mice. The MGMP is part of the Australian Phenomics Network. Excision of exon 5 (ENSMUSE00001068050) of the longest *Axdnd1* transcript ENSMUST00000213088.1 (Supplementary Figure 1; transcript 211) was achieved using CRISPR guide sequences shown in Table 2 and led to a premature stop codon in exon 6. This modification generated two founder lines: one with a 564 bp deletion and 5 bp insertion (*Axdnd1^del1^* line) and another with a 571 bp deletion (*Axdnd1^del2^*line) – both contained a premature stop codon in exon 6. Deletions were validated via Sanger sequencing at Micromon (Monash University, Australia). After mating founder mice to wild type C57BL/6J mice, resulting mutant *Axdnd1^+/-^*mice were used to generate *Axdnd1*^-/-^ individuals. *Axdnd1* expression levels for the longest transcript (*Axdnd1-*211) and a smaller transcript (*Axdnd1-* 208) that does not contain exon 5 (and thus was not targeted by the CRISPR deletion) were assessed in testes from knockout males (*Axdnd1^del1^*) via qPCR as previously described (41), using primers shown in Table 2.

**Table 2.**
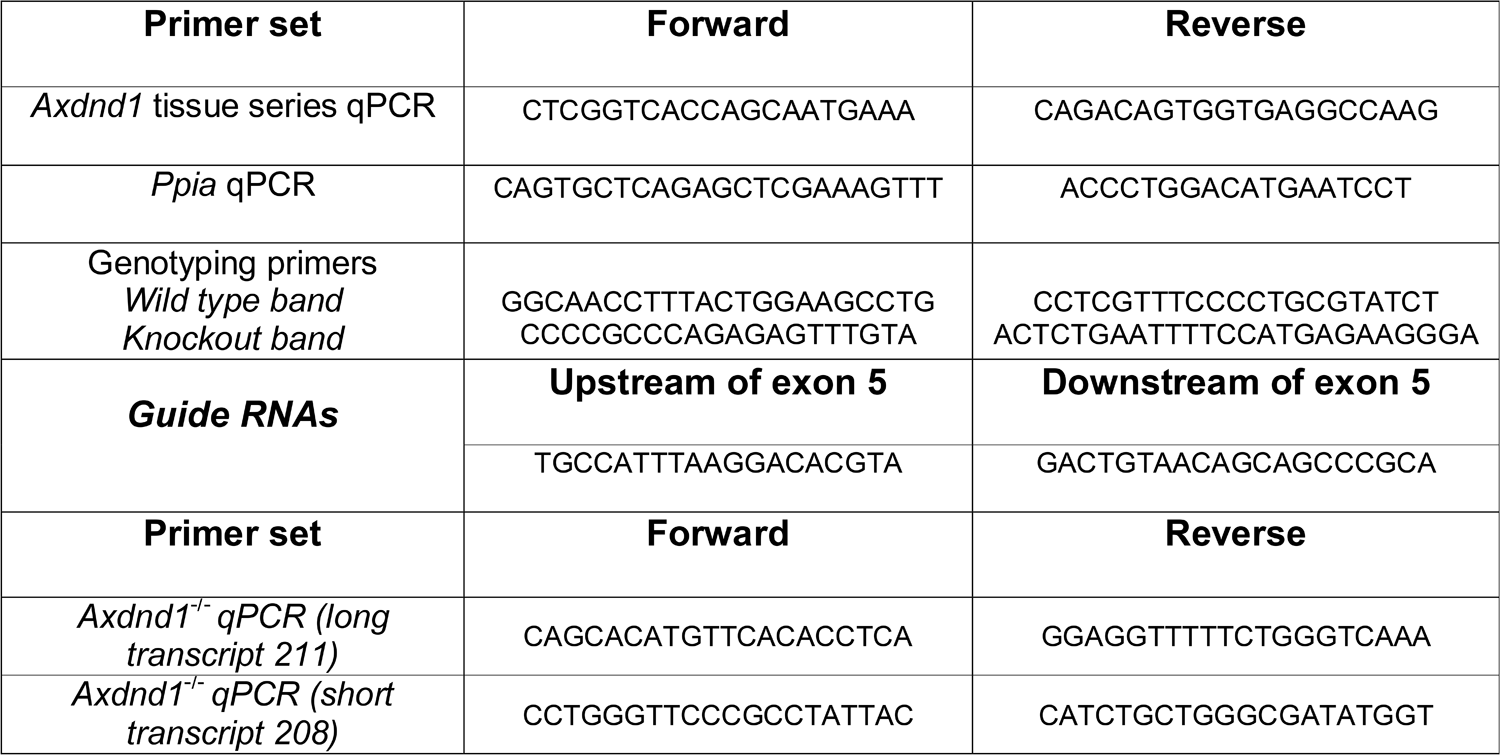
Primer sets used in this study.

### Analysis of *Axdnd1* expression

Whole organ RNA was extracted from adult mouse brain, epididymis, heart, liver, lung, spleen and testis to investigate the expression of *Axdnd1* across different tissues. RNA was also extracted from the testes of mice aged 0-50 days to investigate *Axdnd1* expression throughout the first wave of spermatogenesis. Primers used to detect *Axdnd1* and housekeeping gene *Ppia* are shown in Table 2 below. As a direct measure of *AXDND1* and *Axdnd1* germ cell expression, we also utilised single cell RNA sequencing data generated through previous studies in our lab (24, 25).

### Fertility analysis of *Axdnd1*^-/-^ males

Fertility analyses in this paper were performed on males from the *Axdnd1^del1^* line over an age range of 30-180 days. Phenotyping was also performed on adult males from the *Axdnd1^del2^* line at 70-90 days of age. The fertility and reproductive parameters of wildtype and *Axdnd1*^-/-^ males were assessed using the pipeline outlined in (42). In brief, at least four mice of each genotype were assessed for fertility at two age points (42-50 and 70-80 days) by mating with two young wild type females each. These ages reflect just after the first wave of spermatogenesis and epididymal maturation has completed, and mature adult males, respectively. The presence of a copulatory plug was used as a marker of successful mating. Litter sizes were recorded as the number of pups generated per plug.

At each age point tested, males were culled and weighed, then testes and epididymides were dissected and weighed. One testis was fixed in Bouin’s solution for 5 h and the other was snap frozen on dry ice for calculation of daily sperm production (39) or RNA/protein extraction. One epididymis was fixed in 4% paraformaldehyde for 5 h for histological assessment, while the cauda and vas deferens of the other was used to isolate sperm and any other cells present into MT6 medium using the backflushing method as described previously (41). Sperm motility was assessed via computer assisted sperm analysis as described previously (43), including in an additional experiment where 1 mM of membrane permeable ATP (ATP disodium salt hydrate [A2383 – Merck, Australia]) was added in an attempt to rescue motility. Remaining sperm (and immature germ cells) extracted from the cauda epididymis were washed in 1 × PBS then fixed in 4 % paraformaldehyde for 10 min and resuspended in PBS. Cells were then settled on SuperFrost+ slides overnight at 4 °C for further analysis. In additional experiments, epididymides were collected for calculation of epididymal sperm content (39), and testes and epididymides were collected for and fixed for transmission electron microscopy analysis as detailed below.

### Histological assessment of reproductive tissues and sperm/epididymal luminal cell content

Fixed testes and epididymides were alcohol processed and embedded into paraffin wax using standard methods and 5 µm sections were cut with a microtome. Sections were dewaxed using standard protocols. Testis sections were stained with periodic acid-Schiff’s and haematoxylin reagents (PAS), while epididymis sections were stained with haematoxylin and eosin. Sperm and other cells settled to SuperFrost+ slides were stained with haematoxylin and eosin. All slides were dehydrated in ethanol and xylene baths, then mounted in DPX under a 1.0 × coverslip for light microscope analysis using an Olympus BX-53 microscope fitted with an Olympus 392 DP80 camera.

To quantify the age-related changes in testis histology the incidence Sertoli cell-only tubules, severe hypospermatogenesis and vacuoles, were counted at 40, 70 and 180 days of age, n=3 mice/age/genotype. Sertoli cell-only was assigned when histology showed no germ cells and only Sertoli cells in a tubule, while severe hypospermatogenesis (approaching Sertoli cell-only) was recorded when very few germ cells were present. Intact tubules contained no defects in each of the three categories. Similarly, round spermatid content was determined at postnatal day 22 by counting round spermatid numbers in a minimum of 10 stage V equivalent tubule cross-sections per animal to investigate germ cell commitment to spermatogenesis (n=5/genotype). For this analysis, we focused on round tubules; tubules running longitudinally were excluded.

To quantify apoptosis in the testis of 40-, 70-, and 180-day old males, testis sections were co-stained with 0.5 μg/ml cleaved caspase 3 (Cell Signalling Technology, Australia, Ab 9664) and 0.5 μg/ml cleaved caspase 7 (Cell Signalling Technology, Australia, Ab 9491) antibodies using the immunohistochemical protocol detailed above. The number of caspase positive cells per tubule was then counted in at least 50 tubules chosen randomly per male (n > 3 mice/genotype/age).

To investigate immune cell infiltration, testes were dissected and immersed in Tissue-Tek O.C.T. reagent (Sakura Finetek, CA, USA) prior to freezing in an ethanol dry ice slurry. Frozen testes were cryo-sectioned onto SuperFrost+ slides then stained with CD45 (BD Pharmingen [BD Biosciences] 550539) and 0.5 µg/ml smooth muscle actin primary antibody (Sigma A2547) antibodies overnight at 4 °C, after which they were stained with appropriate secondary antibodies, followed by 10 µg/ml DAPI (Thermo Fisher Scientific).

AXDND1 localisation was investigated in germ cells and sperm via tubule squash preparations from wild type and knockout (negative control) testes using previously established methods (27). Fixed germ cells were incubated in 6 µg/ml AXDND1 antibody (Sigma HPA071114) overnight at 4 °C and stained with appropriate secondary antibodies, 2 µg/ml wheat germ agglutinin-Alexa Fluor 555 (acrosome marker; Thermo Fisher Scientific W32464) and 10 µg/ml DAPI. Attempts to localise AXDND1 via immunohistochemistry and the use of western blotting on whole testis tissue/lysate failed to produce specific staining or bands.

### Investigation of the blood-testis barrier integrity and immune infiltration

A biotin tracer (EZ-Link™ NHS-Biotin; Thermo Fisher Scientific, Australia) was used to test the patency of the blood-testis barrier in *Axdnd1*^-/-^ mice as described previously (44). Briefly, after nicking the capsule, testes were incubated in 10 mg/ml biotin in 0.05 M CaCl_2_ for 30 min, fixed in Bouin’s solution for 5 h and then processed into sections as described above. Sections were dewaxed and prepared for staining as detailed above. Staining included incubation with 0.5 µg/ml smooth muscle actin primary antibody (Sigma A2547) overnight at 4 °C, followed 1 µg/ml donkey anti-mouse secondary and 2 µg/ml streptavidin-Alexafluor-488 reagent (to visualise the biotin, Thermo Fisher Scientific). Nuclei were counterstained with 10 µg/ml DAPI (Thermo Fisher Scientific) for 20 min and slides were mounted with Dako fluorescence mounting medium (Agilent).

### Investigation of spermatogonia dynamics during the first wave of spermatogenesis

To quantify numbers of spermatogonia, including those committing to spermatogenesis, during the first wave of spermatogenesis, stained investigated germ cell content in testis sections at postnatal days 3-7 and 22-35. Testis sections were stained with the germ cell marker MVH (day 3-7; 3 µg/ml, Abcam ab13840), the pro- and undifferentiated spermatogonia marker PLZF (day 5-35; 0.6 µg/ml, R&D Systems AF2944), and dual stained with the transcription factor marker FOXO1 (day 7; 1 µg/ml, Cell Signaling Technology 2880) and differentiating spermatogonia marker c-KIT (day 7; 0.6 µg/ml, R&D Systems AF2944) overnight at 4 °C. Sections were co-stained with 0.5 µg/ml smooth muscle actin antibody (Sigma A2547) and incubated with relevant secondary antibodies then 10 µg/ml DAPI to stain DNA. The number of tubules with positive staining for MVH or PLZF as well as the germ cells positive for each marker (MVH, PLZF, FOXO1 and c-KIT) in at least 20 round tubules was counted (n ≥ 3 animals/genotype). Round spermatid numbers were also counted in at least ten tubules of PAS-stained sections at postnatal day 22 (n = 5 animals/genotype).

### Electron microscopy

To investigate germ cell and sperm ultrastructure, testes were fixed in 5% SuperFix solution (5% glutaraldehyde, 0.2% saturated picric acid and 5% paraformaldehyde in 0.1 M sodium cacodylate) at room temperature for 5 hours, and then overnight at 4 °C. After one hour of fixation, a small incision was made along the tunica albuginea of each testis, and after 5 hours the testes were cut in half to facilitate optimal fixation of tissue. Images were taken on a Jeol 1400 Plus electron microscope at the Vera and Clive Ramaciotti Centre for Electron Microscopy (Monash University, Australia), a Talos L120C or a FEI Teneo VolumeScope at the Ian Holmes Imaging Cente (Bio21 Institute, Parkville, Australia). To view the sperm tail ultrastructure, sperm were isolated from the cauda epididymis and incubated in 100 µl of 1 x PBS for 30 min to strip the plasma membrane as previously described (45). Samples were then diluted in 0.1 M sodium cacodylate buffer and settled onto gold coated glass.

### Statistical analysis

Statistical analyses to determine significance between wild type and *Axdnd1*^-/-^ parameters at each age point were performed in Prism (GraphPad, San Diego, USA) using an unpaired t-test with α = 0.05. For age series data, a t-test was used to compare genotypes at each age point. Significance of apoptosis data was assessed using a script in RStudio as previously described (46).

### Study approval

#### Patient description

As a described previously (20), we used exome sequencing on infertile men presenting with azoospermia or severe oligozoospermia to investigate novel genetic causes of male infertility. Herein, we identified additional variants in *AXDND1*. The variant in patient 1 was annotated, filtered and prioritised as per (20). Variants in patients 2 (M1557) and 3 (M2628) were identified in the Male Reproductive Genomics (MERGE) cohort including 2412 participants (n = 2127 men with severe/extreme oligo-, crypto- or azoospermia) according to (47). Testicular biopsies for histological examination have been taken in parallel with biopsies for therapeutic testicular sperm extraction and processed as described in (48). Experimental ethics for patients 2 and 3 were approved by the Münster Ethics Committees and Institutional Review Boards (2010-578-f).

#### Animal ethics

Experimental procedures involving mice followed animal ethics guidelines generated by the Australian National Health and Medical Research Council (NHMRC). All animal experiments were approved by the Animal Experimentation Ethics Committee (BSCI/2017/31) at Monash University, Clayton or The University of Melbourne Animal Ethics Committee (application 20640).

## Data availability

Data are available upon request to the corresponding author.

## Author contributions

BJH, JM, JN, and AEOC undertook experiments.

BJH analysed the data.

BJH, RH, JD and MKOB designed the experiments.

BJH and MKOB wrote the paper.

The patients were recruited by AL and SK, and their DNA sequenced by LN, CF, FT, KIA and DFC.

All authors provided intellectual input and feedback on drafts.

All authors reviewed and approved the final publication.

## Acknowledgements

Luisa Meyer for technical support.

