## Supplementary figures and images for "AXDND1 is required to balance spermatogonial commitment and for sperm tail formation in mice and humans"

### Supplementary Figure 1

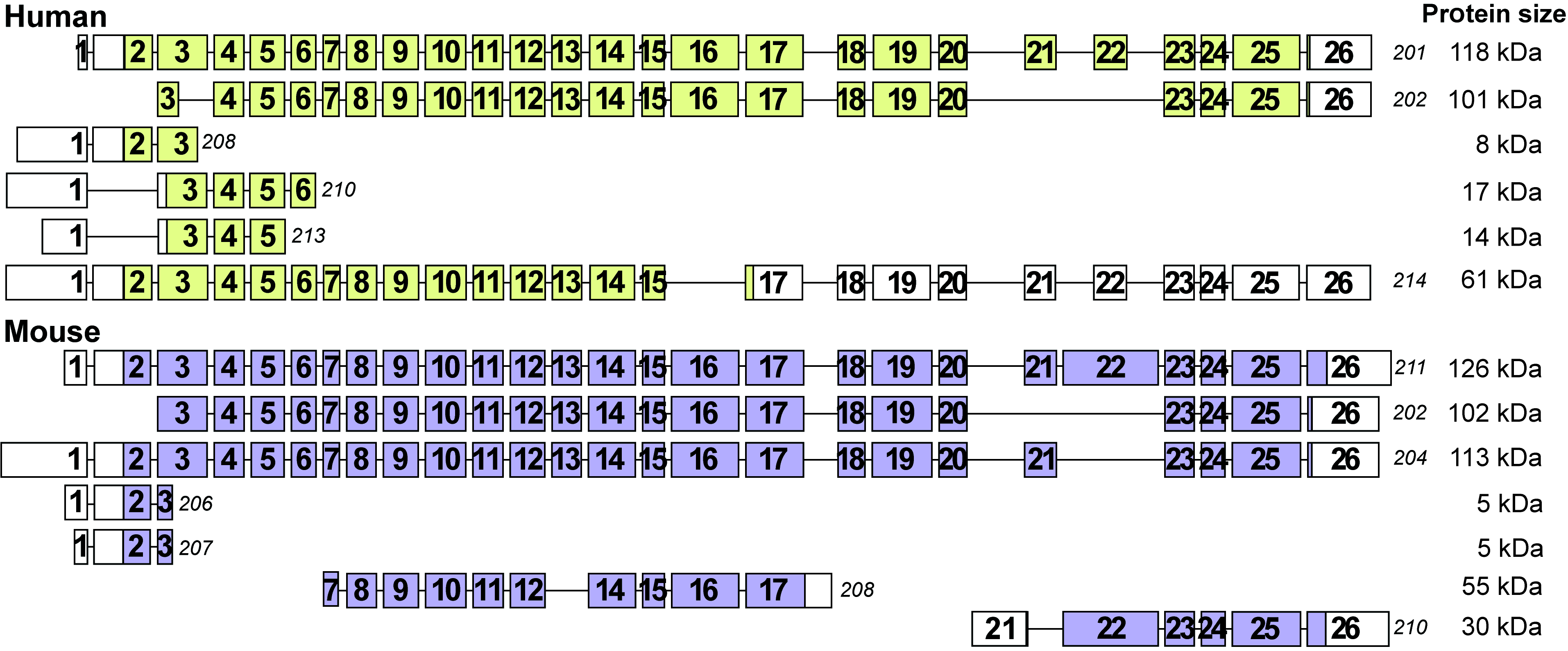

### Supplementary Figure 2

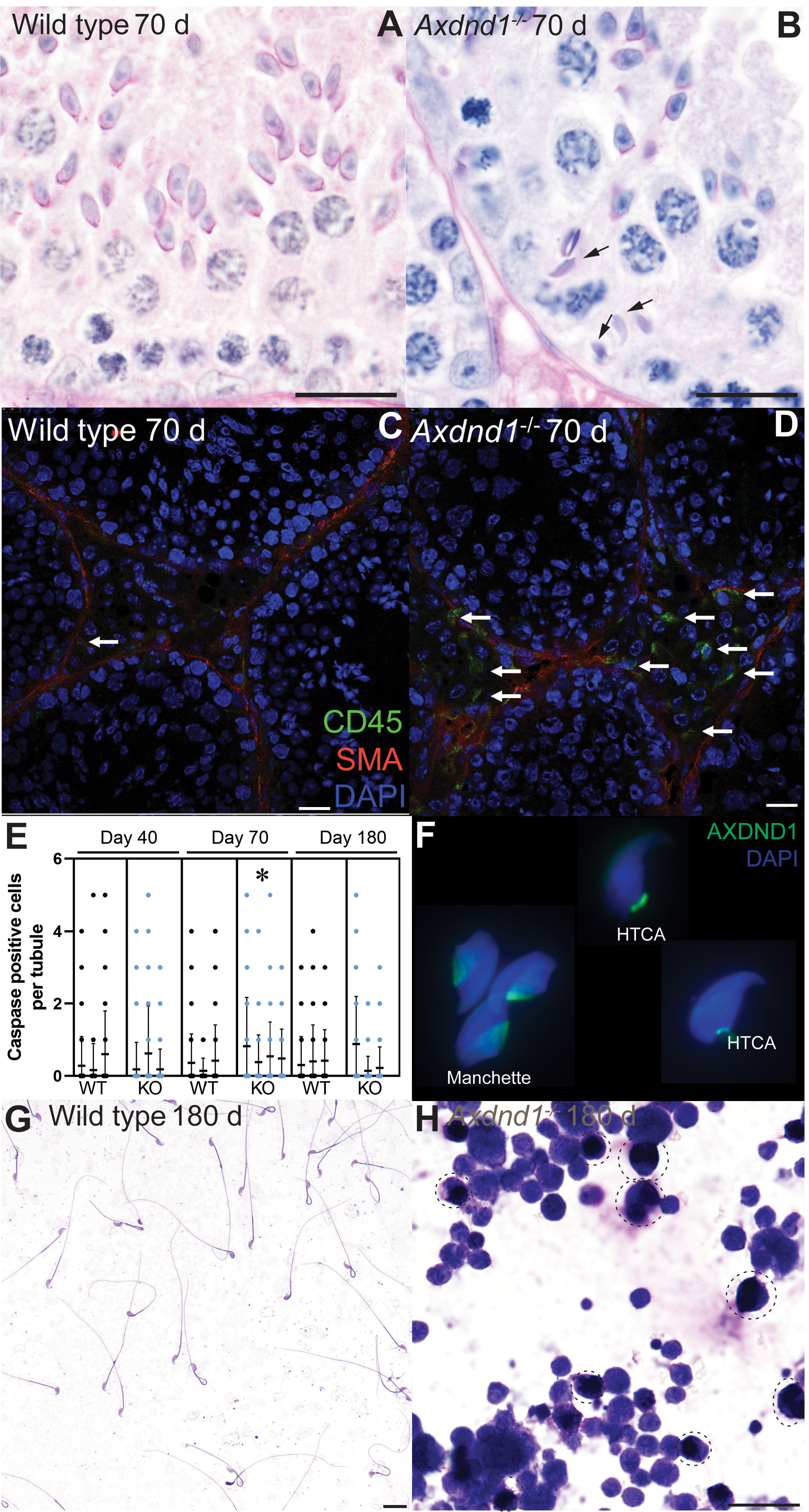

### Supplementary Figure 3

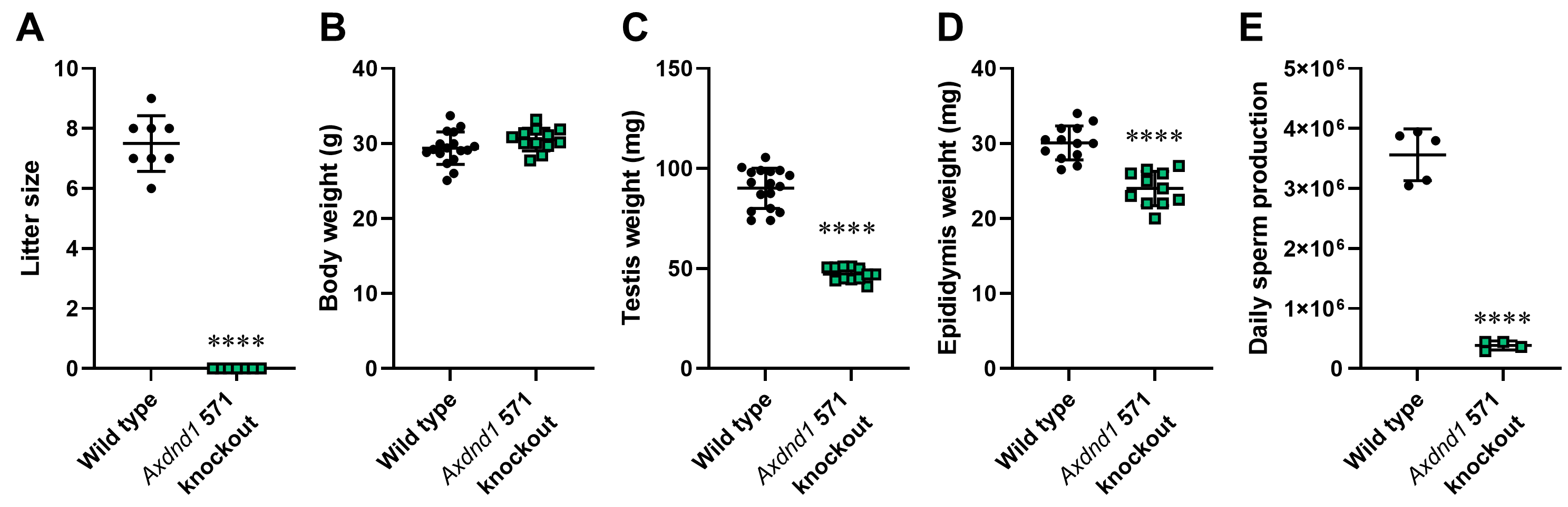

### Supplementary Figure 4

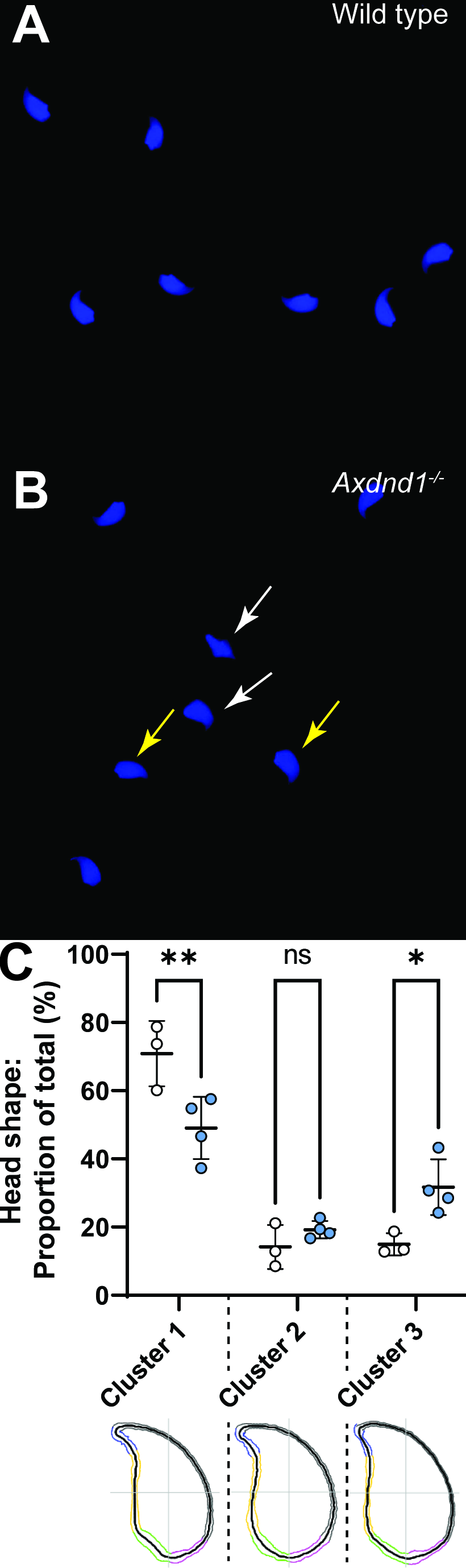

### Supplementary Figure 5

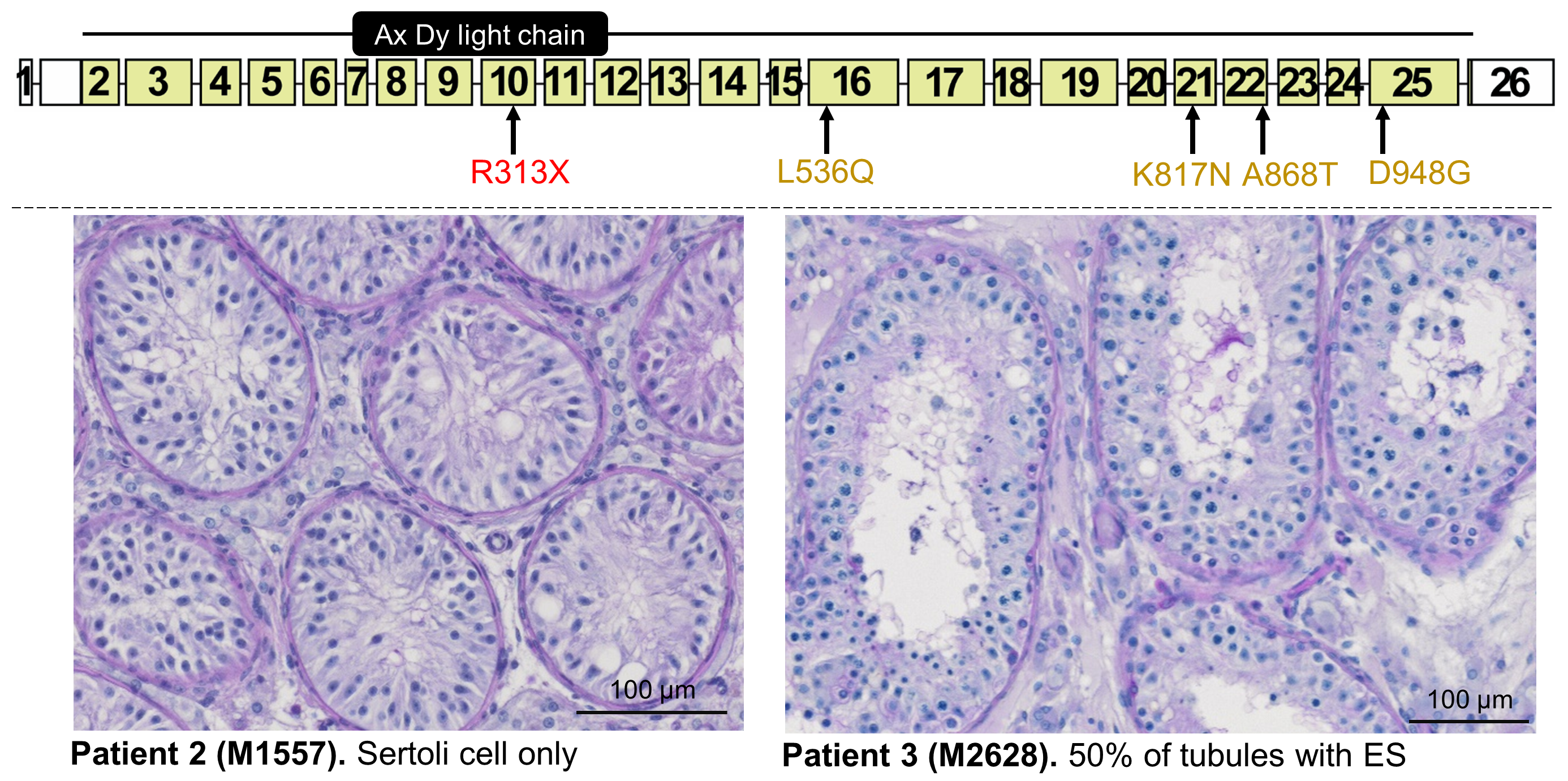
